# Effect of spatiotemporally changing environment on serial dependence in ensemble representations

**DOI:** 10.1101/2021.11.30.470662

**Authors:** Sangkyu Son, Joonyeol Lee, Oh-Sang Kwon, Yee-Joon Kim

## Abstract

The recent visual past has a strong impact on our current perception. Recent studies of serial dependence in perception show that low-level adaptation repels our current perception away from previous stimuli^1–5^ whereas post-perceptual decision attracts perceptual report toward the immediate past^6–12^. In their studies, these repulsive and attractive biases were observed with different task demands perturbing ongoing sequential process. Therefore, it is unclear whether the opposite biases arise naturally in navigating complex real-life environments. Here we only manipulated the environmental statistics to characterize how serially dependent perceptual decisions unfold in spatiotemporally changing visual environments. During sequential mean orientation adjustment task on the array of Gabor patches, we found that the repulsion effect dominated only when ensemble variance increased across consecutive trials whereas the attraction effect prevailed when ensemble variance decreased or remained the same. The observed attractive bias by high-to-low-variance stimuli and repulsive bias by low-to-high-variance stimuli were reinforced by the repeated exposure to the low and the high ensemble variance, respectively. Further, this variance-dependent differential pattern of serial dependence in ensemble representation remained the same regardless of whether observers had a prior knowledge of environmental statistics or not. We used a Bayesian observer model constrained by visual adaptation^13,14^ to provide a unifying account of both attractive and repulsive bias in perception. Our results establish that the temporal integration and segregation of visual information is flexibly adjusted through variance adaptation.

## Results

### Previous perception integrates with current perception when environmental variance remains the same

Our first aim was to investigate whether current ensemble representation is biased toward previous ensemble representation when ensemble variance does not change. To test for serial dependence in ensemble representations, observers completed a series of trials in which they were presented with the array of sixteen tilted Gabor patches in the peripheral visual field. The standard deviation of the sixteen Gabors’ orientations was either 5° (low-variance; L) or 20° (high-variance: H) on each trial, and the mean was randomly selected from -40° to 40°, in steps of 10° relative to the previous mean orientation. By manipulating the transition probability between low-variance and high-variance trial (Figure 1B), we exposed observers to either a repetitive (Figure 1B, the 1^st^ column) or a random environment (Figure 1B, the 2^nd^ column). Observers were asked to report the perceived mean orientation of each Gabor array by adjusting a response bar (Figure 1A). We then analyzed the dependence of the adjustment responses on the mean orientation of the previous trial for the possible pairs of consecutive trials. When the consecutive trials had the same level of orientation variability in their Gabor arrays (e.g. LL and HH), we found that adjustment responses were systematically attracted toward the mean orientation of the previous trial regardless of whether or not these consecutive trials were extracted from repetitive (Figure 2A) or random (The 1^st^ and 2^nd^ columns in Figure 3A) session.

**Figure 1.**
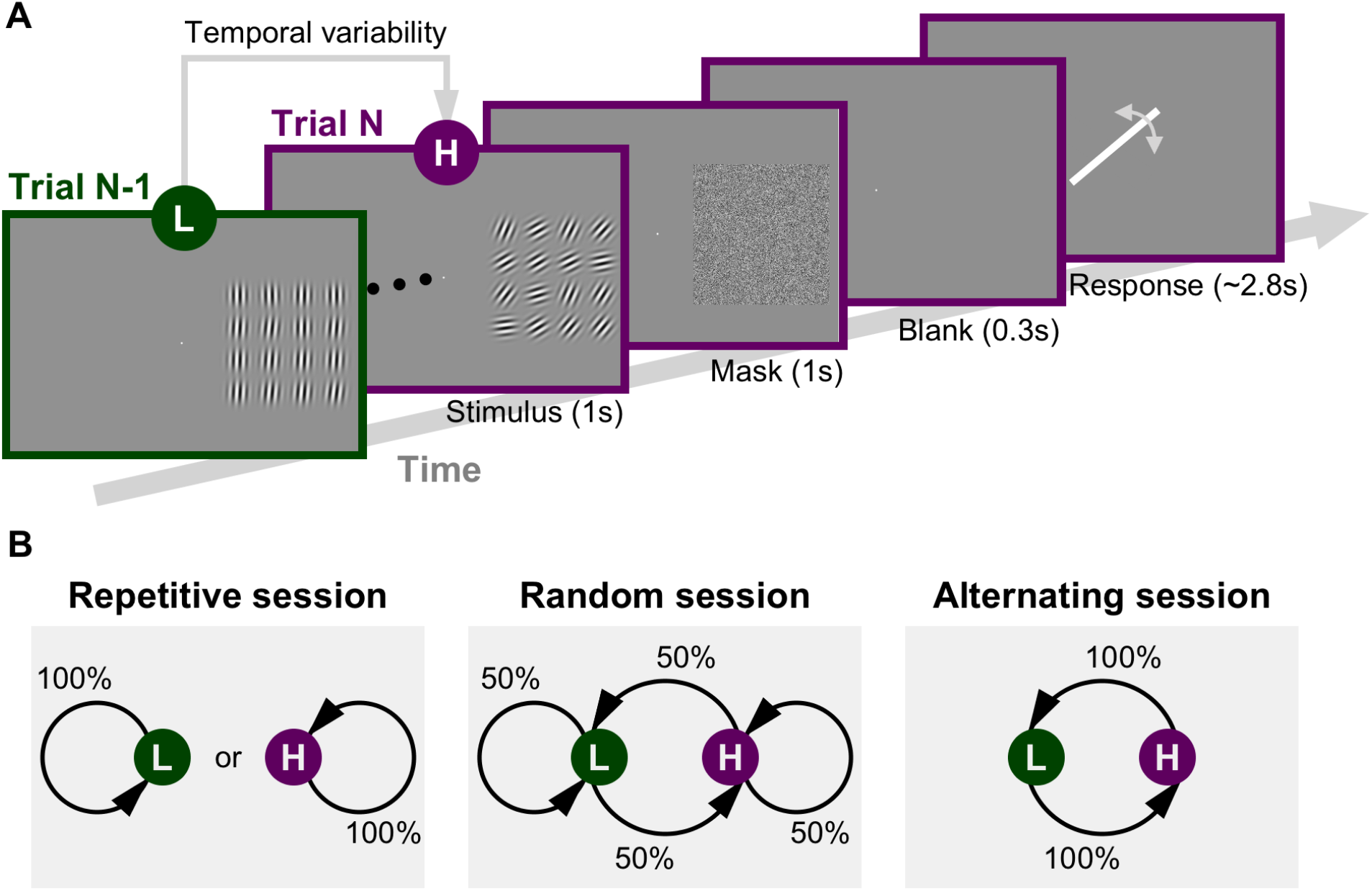
Schematic trial sequence and experimental sessions. (A) On each trial, a 4 × 4 array of sixteen Gabors was peripherally presented for 1000 ms, followed by a 1000 ms salt and pepper noise mask to reduce afterimages, and a 300 ms blank. Observers were instructed to match an orientation bar to the perceived mean orientation of the Gabor array by adjusting the orientation of a response bar. The tilt of each Gabor patch was drawn from a uniform distribution with the low (L, s.d. = 5°) or the high (H, s.d. = 20°) standard deviation. An example trial sequence of L trial *N-1* and H trial *N* (LH) was shown. (B) Three transition probabilities between the low-variance array and the high-variance array of the consecutive trials yielded three environmental sessions: Repetitive (transition probability of 0), Random (transition probability of 0.5) and Alternating (transition probability of 1). Observers were informed of the environmental statistics in alternating session but not in random session and repetitive session.

**Figure 2.**
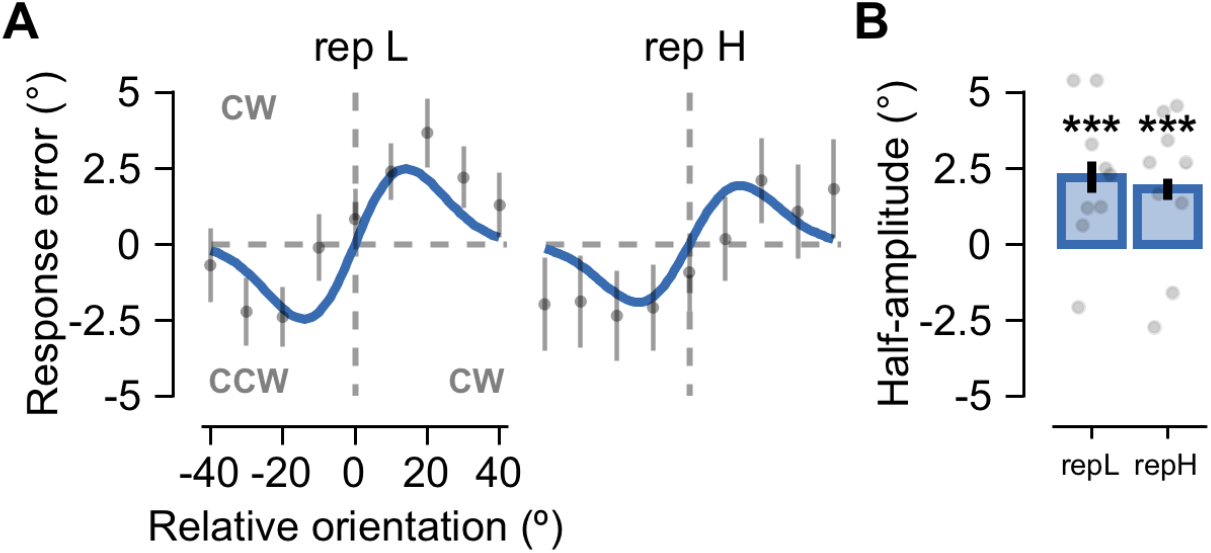
Positive serial dependence under repetitive environment. (A) Serial dependence error plots of all trials in the two repetitive trial combinations of the low-variance array (rep L) and the high-variance array (rep H), respectively. The x-axis is the previous mean orientation minus the current mean orientation. The y-axis is response bar orientation minus the mean orientation on the current trial (i.e. an error in the adjustment task). The blue lines show a derivate-of-Gaussian (DoG) curve fit to the average response error (gray dots). Error bars indicate bootstrapped 95% confidence intervals. In both rep L and rep H experimental conditions, the DoG curves show that the response errors are biased toward the previous mean orientation. (B) In order to quantify the magnitude of serial dependence, the mean bootstrapped half-amplitude of the DoG curve was calculated for each observer (see Methods for details). The left and right blue bars and gray dots represent the averaged and individual mean bootstrapped half-amplitudes for rep L and rep H experimental condition, respectively. Observers showed significant positive serial dependence in both combinations in the perceived mean orientation of current Gabor array (permutation test, *ps* < 0.001). Error bars indicate bootstrapped 95% confidence intervals.

**Figure 3.**
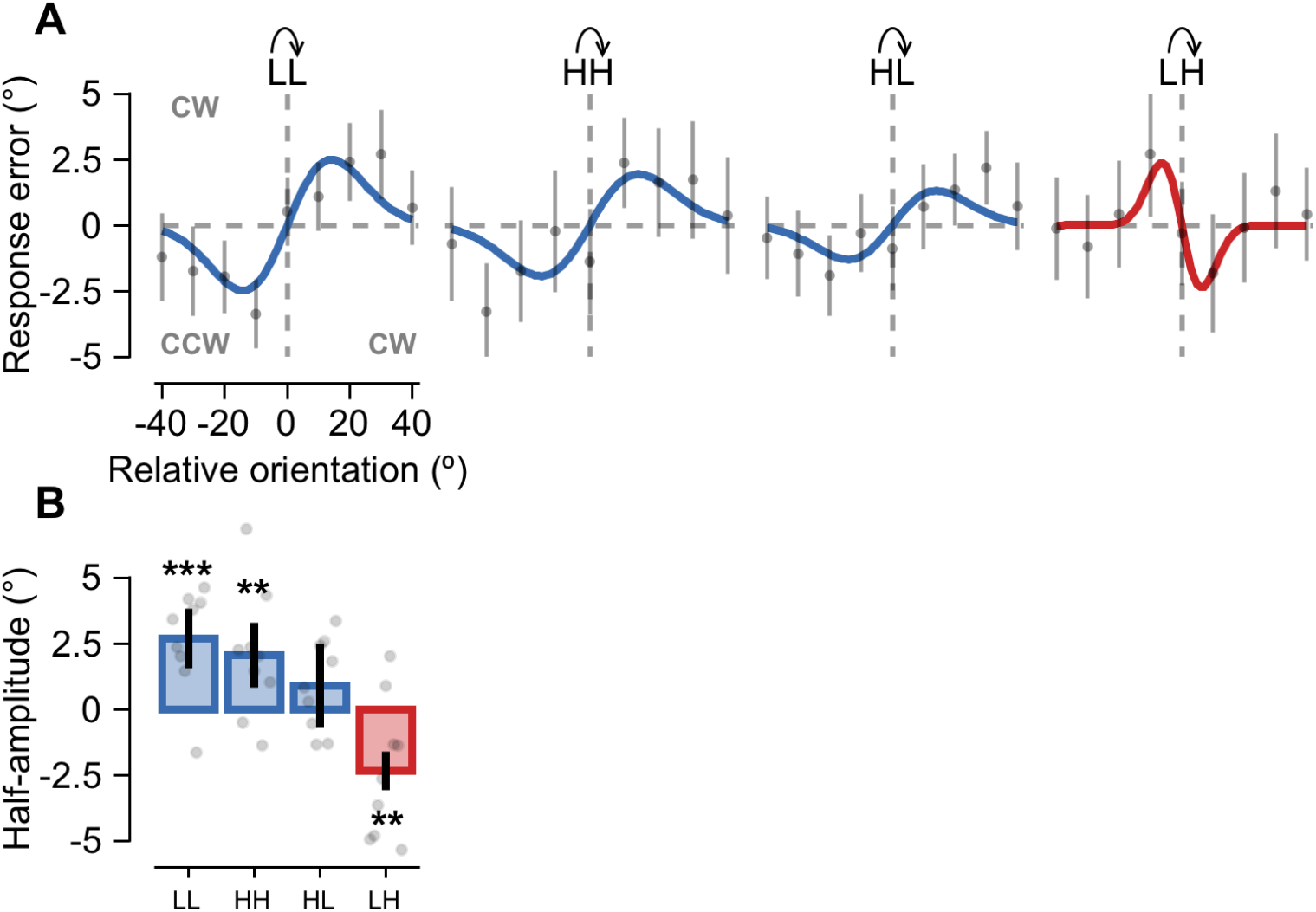
Positive and negative serial dependence under random environment. (A) Serial dependence error plots of current trials (gray dots for average response errors and error bars for bootstrapped 95% confidence intervals) in the four consecutive trial combinations of the low-variance and the high-variance arrays (LL, HH, HL, and LH). Three blue and a red line show DoG curve fits to the data from LH, HH, HL, and LH, respectively. Black arrows above the combination label indicate the trials the serial dependence is computed from. (B) Serial dependence magnitude of the previous trial was computed for each consecutive trial combination (LH, HH, HL, and LH). The mean bootstrapped half-amplitudes showed significant positive serial dependence when ensemble variance does not change over the consecutive trials (LL and HH), and no specific biases when ensemble variance decreases in time (HL), while evident negative serial dependence when ensemble variance increases in time (LH) (Half-amplitude permutation test, p_LL_ < 0.001 p_HH_ = 0.008, p_HL_ = 0.115, p_LH_ = 0.003, see Methods for details). Bar and gray dots indicate the average and individual observers, respectively, and error bars indicate bootstrapped 95% confidence intervals.

As a measure of the serial dependence, we computed the half-amplitude of a simplified derivative-of-Gaussian (DoG) that was fit to each observer’s data (See Methods for the details). We then bootstrapped each observer’s data with 5000 iterations and reported the mean bootstrapped half-amplitude (Figures 2B and 3B). For the pairs of consecutive trials with the same orientation variability, all observers displayed a significantly positive DoG half-amplitudes, indicating that perceived mean orientation on a given trial was significantly pulled in the direction of the mean orientation presented in the previous trial (Group-level permutation test, Figure 2B: *p*_*rep L*_ < 0.001, *p*_*rep H*_ < 0.001 based on group permuted null and the two leftmost bars in Figure 3B: *p*_*LL*_ < 0.001, *p*_*HH*_ = 0.008 based on group permuted null). These positive biases toward the previous mean orientation in the trial combinations of LL and HH even from random environment where ensemble variance was constantly changing across trials strongly suggest that the repetition of the adapted ensemble variance rapidly facilitates visual stability. Especially the positive bias in the trial combination of HH extended the previous finding of positive serial dependence in the consecutive trials with low ensemble variance^12^. This robust and easily manifesting attribute of the positive bias suggests that the vision essentially integrates representations over time through temporal variability adaptation for the sake of stability.

### Current perception repels away from previous perception when environmental variance increases over time

Next, we examined how the change in ensemble variance influenced on serial dependence in ensemble representations. In the random session, observers did not show any prominent serial dependence bias when the ensemble variance decreased across consecutive trials (HL: The 3^rd^ column in Figure 3A). Observers’ adjustment responses, however, were systematically repelled away from the mean orientation of the previous trial when the ensemble variance increased across trials (LH: The 4^th^ column in Figure 3A). The DoG curve fit of the response error data showed that observers displayed a significantly negative half-amplitude when the ensemble variance increased across consecutive trials and no systematic bias when the ensemble variance decreased across successive trials (Group-level permutation test, Figure 3B; *p*_*LH*_ = 0.035 based on group permuted null, *p*_*HL*_ = 0.176 based on group permuted null).

To investigate whether the negative repulsive bias in the trial combination of LH was observed by chance or not in the random session, we generated the null distribution for each trial combination by randomly regrouping the mean bootstrapped half-amplitudes of all observers into four trial combinations of LL, HH, HL, and LH with 10,000 iterations (See Methods for the details). By comparing the observed half-amplitude with the null distribution in each trial combination, we found that the serial dependence in trial combination of LH was qualitatively different from the serial dependence in the other three trial combinations that similarly showed the positive bias (*p*_*LH*_ < 0.001, corrected).

However, it is still possible that the positive and the negative serial dependence could be due to motor response bias^15^, oblique bias^7^, or localization bias^16^. To control for these potential confounds, we performed another statistical test on the response error data. We randomly shuffled trial orders and generated the null distribution of half-amplitudes for each rearranged trial combination of each observer with 10000 iterations. We then examined if the observed half-amplitude was significantly different from the mean half-amplitude of the null distribution for each trial combination. The serial dependence in the four trial combinations remained the same as observed in Figure 3 (*t*-test, *p*_*LL*_ = 0.003, *p*_*HH*_ = 0.034, *p*_*HL*_ = 0.183, *p*_*LH*_ = 0.039).

Overall, these analyses clearly show that when observers perform exactly the same orientation adjustment task, their adjustment responses are repelled away from the previous mean orientation when ensemble variance increases, but are attracted toward the previous mean orientation in most environmental changes. These results suggest that the changing environmental statistics are the main cues for either integration or differentiation between successive representations. Specifically, increased spatial variability initiates the control process to segregate the current and previous information to promote change detection while unchanged or decreased spatial variability initiates the control process to integrate the current and previous information to support visual stability over time. Therefore, the ultimate consequence of serial dependence depends on the spatiotemporal variability that adaptively controls the balance of attractive and repulsive effect.

### Repulsive perceptual bias for current high-variance stimulus increases with repeated exposure to low-variance environment

Despite the same amount of change in spatiotemporal variability, we wondered why serial dependence differs depending on whether ensemble variance increases (LH in Figure 3) or decreases (HL in Figure 3) over successive trials. Based on the previous findings that the highly heterogeneous array stimulus degrades the accuracy of ensemble representation [22–25] and the innately volatile feature such as facial expression induces repulsive bias [15], we hypothesize that much weaker low-level orientation specific adaptation to the high-variance stimulus than low-variance stimulus differentially modulates serial dependence by the high-to-low-variance and the low-to-high-variance trials. If spatiotemporal variability is a cue for differentiation or integration process, our prediction is that low-level sensory adaptation would work to tip the balance of repulsive and attractive effect in favor of differentiation process when ensemble variance increases across successive trials. Therefore, given the fact that the temporally constant spatial variability (LL and HH, Figure 3) fundamentally involves a representation of the current stimulus biased toward a representation of previous stimulus, differential adaptation effect tips the balance between perceptual memory demands in favor of repulsion or attraction depending on the history. When high-variance trials continue, positive bias accumulates and then the accumulated positive bias remains when low-variance trial appears because high-variance stimulus has a weak adaptation effect on the low-variance stimulus. On the other hand, when low-variance trials continue, positive bias again accumulates and then repulsive bias will be bigger because low-variance stimulus has strong adaptation effect on the current high-variance stimulus. To verify this hypothesis, we sorted trials into two groups based on the repetition number of low-variance trials presented before high-variance trial (e.g. LH, LLH, and LLLH) and the repetition number of high-variance trials before low-variance trial (e.g. HL, HHL, and HHHL). We separately performed linear trend analysis on half-amplitudes of DoG curves in each group of the sorted trials.

We found that the adjustment response was increasingly more repelled away from the previous mean orientation as more low-variance trials were repeatedly presented before the current high-variance trial (Figure 4A). The half-amplitudes of DoG curves for the successive trials of low- and high-variance Gabor array showed a negatively increasing trend as a function of the repetition number of low-variance trials (*p*_*linear trend*_ = 0.042; *p*_*LH*_ = 0.035, *p*_*LLH*_ = 0.006, *p*_*LLLH*_ < 0.001, Figure 4B). On the other hand, we found that the more repeated exposure to the high-variance trial before the low-variance trial, the more the observers’ adjustment response was attracted toward the previous mean orientation although each half-amplitude of DoG curve did not show a significant positive bias (Figure 4C-D). These results clearly show that the strength of variance adaptation modulates sensitivity to change, which ultimately determines the fate of serial dependence. The fact that strong adaptation in the context of temporally increased spatial variability under constantly changing environment overrides dominant integration processes provides the evidence for the context-dependent flexibility of serial dependence.

**Figure 4.**
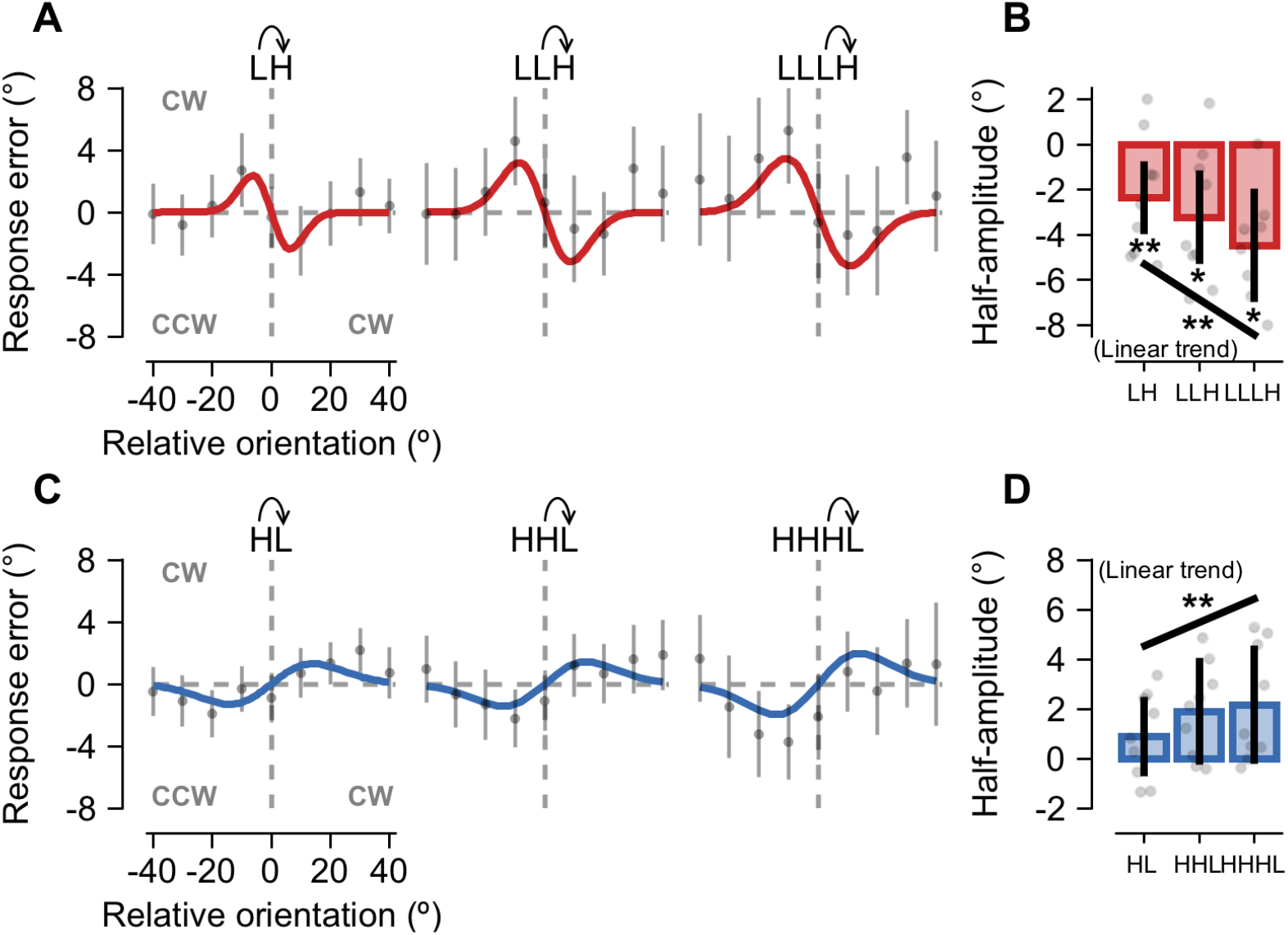
The effect of stationary history before the temporal change of variance on serial dependence. (A) Serial dependence error plots of trials with the high-variance array (gray dots) as a function of the relative mean orientation of the low-variance array presented once, twice, and three times prior to the current trial (LH, LLH, LLLH). Red lines from the 1^st^ column to the 3^rd^ column show DoG curve fits to the data from LH, LLH, and LLLH, respectively. As low-variance array was repeated more beforehand, the DoG curves show that the response errors of current H trials are repelled more away from the previous mean orientation of the low-variance array. Error bars indicate bootstrapped 95% confidence intervals. Black arrows above the combination label indicate the trials the serial dependence is computed from. (B) The mean bootstrapped half-amplitudes of DoG curves computed for LH, LLH, and LLLH (red bars). Linear trend analysis showed that the negative serial dependence magnitudes have a linear trend as the adaptation to the low-variance array gets longer (trend analysis, F = 9.685, p = 0.007). The black line below three bars indicates the linear trend that was tested. Bar and gray dots indicate the average and individual observers, respectively, and error bars indicate bootstrapped 95% confidence intervals. Asterisks indicates whether the trend was significant. (C) Serial dependence error plots of trials with the low-variance array (gray dots) as a function of the relative mean orientation of the high-variance array presented once, twice, and three times prior to the current trial (HL, HHL, HHHL). Blue lines from the 1^st^ column to the 3^rd^ column show DoG curve fits to the data from HL, HHL, and HHHL, respectively. Error bars indicate bootstrapped 95% confidence intervals. Black arrows above the combination label indicate the trials the serial dependence is computed from. (D) The mean bootstrapped half-amplitudes of DoG curves computed for HL, HHL, HHHL (blue bars). Bar and gray dots indicate the average and individual observers, respectively. Linear trend analysis showed that the positive serial dependence magnitudes have a linear trend as the adaptation to the high-variance array gets longer (trend analysis, F = 9.719, p = 0.007). The black line below three bars indicates the linear trend that was tested. Other notations are identical to the (B).

### Prior knowledge of environmental statistics does not alter the flexible interplay between variance adaptation and positive dependency

Given that only the temporally increased spatial variability promotes change detection whereas the temporally decreased or constant spatial variability supports visual stability over time, it is plausible that observers least anticipate the transition from the low- to high-variance trial under constantly changing environment. Thus, we tested if the prior knowledge of environmental statistics can alter the negative repulsive bias in the consecutive trials of the low- and high-variance Gabor array. For this purpose, nine additional observers performed the same mean orientation adjustment task in a session where the low-variance and the high-variance Gabor arrays were alternately presented (The 3^rd^ column in Figure 1B). Importantly, before the whole alternating session started, we explicitly informed observers of the transition probability between the low- and high-variance trial by running several practice trials.

Even though observers knew how environment changes in advance, the pattern of serial dependence in alternating session remained the same as observed in the consecutive trials with different ensemble variance in random session. The temporally decreased spatial variability induced a positive serial dependence and the temporally increased spatial variability induced a negative serial dependence (Figure 5A). The DoG curve fit of the response error data also showed that observers displayed a significantly positive half-amplitude when ensemble variance decreases across the consecutive trials and a significantly negative half-amplitude when ensemble variance increases across the successive trials (Group-level permutation test, Figure 5B; *p*_*altHL*_ = 0.002 based on group permuted null, *p*_*altLH*_ = 0.006 based on group permuted null). These results provide strong evidence that variance adaptation flexibly interacts with dominant positive dependency on the fly depending on spatiotemporal change itself, not on top-down anticipatory prior knowledge of it.

**Figure 5.**
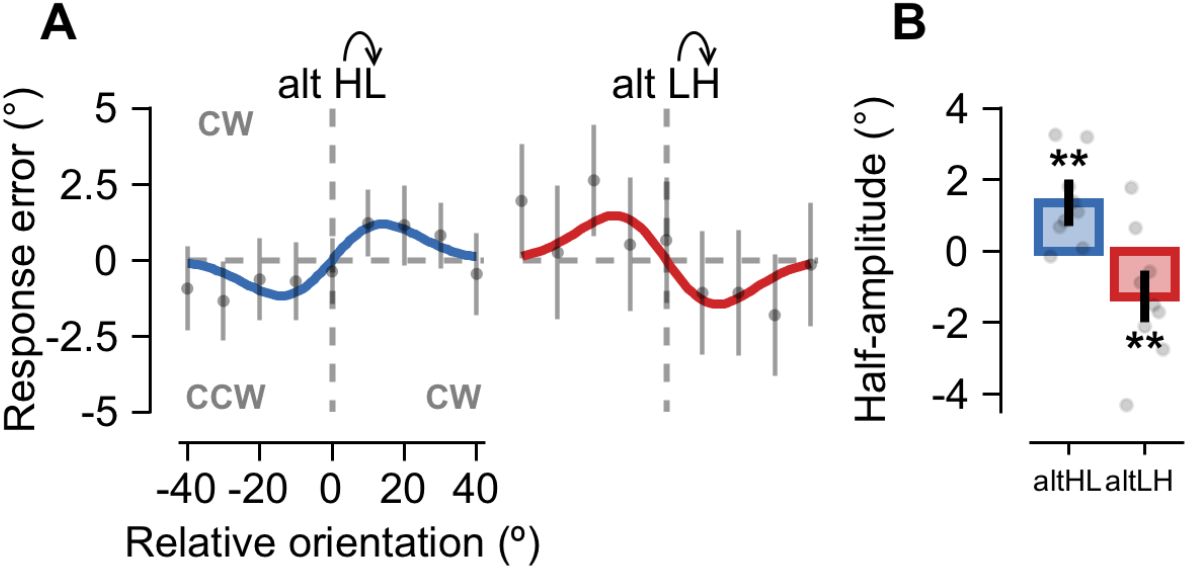
Positive and negative serial dependence under alternating environment. Observers were explicitly informed of the transition probability between L and H trials before the start of the alternating session. (A) Serial dependence error plots of current trials (gray dots) as a function of the relative mean orientation of the previous trial in the two alternating trial combinations of the low-variance and the high-variance arrays (alt HL and alt LH). Error bars indicate bootstrapped 95% confidence intervals. Blue and red lines show DoG curve fits to the data from alt HL and alt LH, respectively. Black arrows above the combination label indicate the trials the serial dependence is computed from. (B) The mean bootstrapped half-amplitudes of DoG curves computed for alt HL (blue) and alt LH (red). The mean bootstrapped half-amplitudes showed significant positive serial dependence in alt HL, but negative serial dependence in alt LH (Half-amplitude permutation test, p_alt HL_ = 0.002 p_alt LH_ = 0.006). Bar and gray dots indicate the average and individual observers, respectively, and error bars indicate bootstrapped 95% confidence intervals.

### An adapted Bayesian observer model captures the context-dependent nature of serial dependence

Recently, the positive recency bias was well explained by a conventional Bayesian observer model that combines the prior expectations about the mean orientation of the Gabor arrays with the immediate likelihood function of the sensory evidence on a given trial^17^. Since our results clearly show that the visual system flexibly controls the interaction between sensory adaptation and positive dependency depending on changes in the environmental statistics, we attempted to better explain both the observed positive and negative serial dependence using an adapted Bayesian observer model^13,14^. This adapted Bayesian model assumes that the sensory adaptation increases the signal-to-noise ratio (SNR) of the measurement around the parameter value in the vicinity of the adaptor (Figure 6A) and decreases the SNR of the measurement further away from the adaptor^18^. This uneven encoding efficiency eventually results in an asymmetric change in the likelihood function (Green solid line in Figure 6B) whose mean is pushed away from the adaptor. This skewed encoding is in agreement with a repulsive effect on the estimate (Red arrow in Figure 6B). In addition, because serially dependent perceptual decision mostly relies on the previous stimulus^6,19–21^, we also hypothesize that on a given trial in a volatile environment, observers have a prior as a weighted mixture of a Gaussian distribution around the mean orientation and a uniform distribution in the range of the mean orientation values that was expected to take outside the previous mean orientation (see Methods for the details, Figure 6C). According to the Bayes rule, the posterior probability distribution is determined by the product of the likelihood function and prior probability distribution (see Methods for the details).

**Figure 6.**
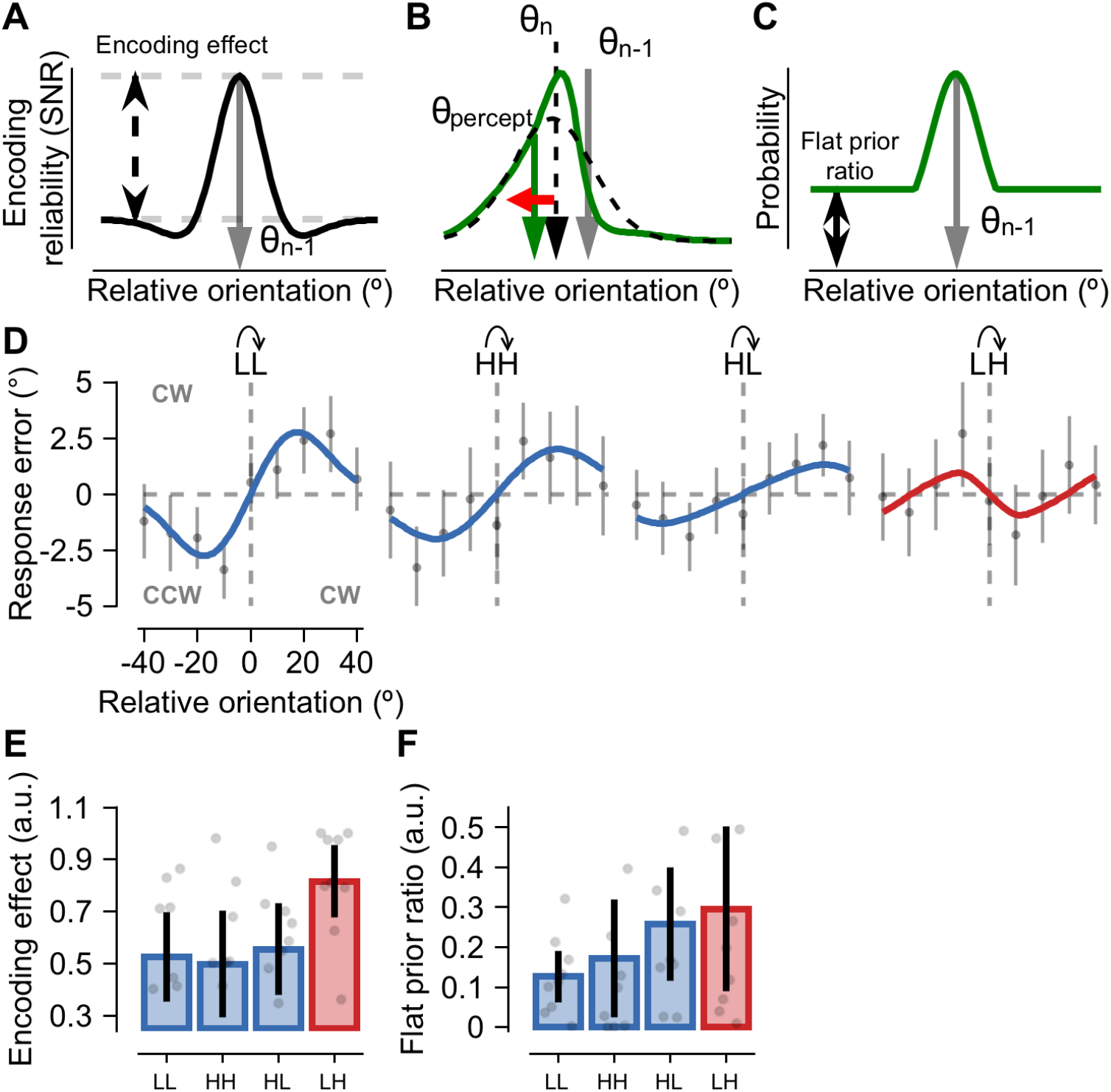
Bayesian observer model with two additional constraints. (A) We assume that adaptation increases the encoding reliability of measurement (SNR) in the vicinity of the orientation of the previous trial n-1^th^ (gray arrow). Encoding effect is the relative height of the encoding reliability. (B) The unevenness in encoding reliability yields an asymmetric likelihood function (green line) as a function of the relative orientation of the previous trial (the gray arrow indicates the given previous orientation), compared to symmetric likelihood function when evenness in encoding reliability was assumed (black dashed line). As a result, the mean of likelihood (green arrow) can be repulsive (red arrow) relatively to orientation currently given (black dashed arrow). (C) The prior belief is assumed to be in Gaussian mixture distribution. The flat prior ratio indicates whether observers prone to rely on prior knowledge or perception. (D) Bayesian model-fitting (red and blue lines) on serial dependence error plots of current trials (gray dots) in the four consecutive trial combinations of the low-variance and the high-variance arrays (LL, HH, HL, and LH). Bayesian model constrained by efficient encoding and Gaussian mixture prior captures not only attractive biases but also repulsive one in serial dependence. Error bars indicate bootstrapped 95% intervals. (E) Encoding effect in LL, HH, HL, and LH combination. In LH, the encoding effect was significantly higher than permutated effect. Shaded region indicates ±2 SEM of permutated distribution. (F) Flat prior ratio in LL, HH, HL, and LH combination. Flat prior did not differ among combinations. Shaded region indicates ±2 SEM of permutated distribution.

We fit this adapted Bayesian observer model to the average adjustment response errors for each pair of four different consecutive trials (Figure 6D). The model successfully captured both the positive and the negative serial dependence depending on the change of environmental statistics (colored lines in Figure 6D). When ensemble variance increases over successive trials, the mean orientation of the previous low-variance stimulus was reliably encoded to have the significant repulsive adaptation effect on estimating the mean orientation of the current high-variance stimulus (Group level permutation test, Figure 6E, p_LL_ = 0.039, p_HH_ = 0.039, p_HL_ = 0.039, p_LH_ = 0.039 based on group permuted null). It is likely that the increased noise in the representation of high-variance stimulus leads to a more asymmetric likelihood function and thus increases likelihood repulsion. The flat prior ratios, however, were not significantly different among the four combinations of consecutive trials although the average flat prior ratio was small when ensemble variance remained the same (Group level permutation test, Figure 6F, p_LL_ = 0.039, p_HH_ = 0.039, p_HL_ = 0.039, p_LH_ = 0.039 based on group permuted null). These results suggest that observers just rely on the previous stimulus rather than acquire a prior over long stimulus history when the to-be-estimated ensemble mean orientation is constantly changing. Further, the fact that the skewed likelihood function depends on the direction of the change in spatiotemporal variability provides the evidence that the sensory adaptation plays a critical role in modulating serial dependence.

## Discussion

In the present study using psychophysical experiments and computational modeling, we show that the waxing and waning of ensemble variance is a crucial cue for biasing the representation of visual stimulus toward or away from the past during serially dependent perceptual decisions on ensemble representations. In a set of three sessions of different environmental statistics, which measure the perceptual estimation of ensemble mean orientation, we found that the temporal integration between current and previous representation, or positive serial dependence, dominates whenever ensemble variance decreases or remains the same across consecutive trials whereas the temporal segregation between current and previous representation, or negative serial dependence, dominates whenever ensemble variance increases over successive trials. We also revealed that the positive attraction effect and negative repulsion effect, observed in the respective case of decreasing and increasing ensemble variance, were reinforced by the repeated exposure to the high and the low ensemble variance, respectively. Moreover, in alternating session we demonstrated that the two opposite biases observed under changes in the variance of visual inputs remained the same regardless of whether observers had a prior knowledge of environmental statistics or not, suggesting variance adaptation as a potential underlying source of automatic switching between integration and differentiation process during serial dependence. Further, based on a previously developed Bayesian observer model constrained by sensory adaptation that elicits the uneven encoding of sensory information in the vicinity of the adaptor^13,14^, we provided a unified account of the environmental statistics-dependent serial dependence by confirming that adaptation has a differential effect on the sensory encoding depending on the history of ensemble variance. This mechanistic switching between attraction and repulsion bias during the sequential mean orientation adjustment task reveals the importance of the environment’s spatiotemporal variability for a complete characterization of the real-time serial dependence.

Our results have a number of implications for understanding the adaptive nature of serial dependence. First, our study directly provides strong empirical evidence that the changes in environmental statistics underlie the flexible switching between attractive and repulsive bias during serially dependent perceptual decisions. Previous studies also reported the existence of the two opposite bias effects of recent history on perceptual and post-perceptual decision processes^1–5,9,22,23^. In these studies, however, the adaptive nature of serial dependence could not be addressed due to the fact that the ongoing temporal dynamics of serial dependence was dissected by using different task structures, different stimulus features, or different response intervals. For example, recent studies used the two separate “adjustment response” and “no adjustment response” or “two-alternative forced-choice perceptual comparison” trials in order to probe the underlying sources of two opposite biases^1,4^. They found that repulsive effect on perception, resembling low-level adaptation, dominate after stimuli that require no perceptual adjustment response and attractive effect on post-perceptual decision emerges during adjustment response condition. Because observers in these studies were cued differently for “adjustment response” trials and “no response” or “forced-choice response” trials, it is possible that different task demands may alter their decisional strategies and hence sequential effects. For this reason, it difficult to clarify whether the switch between positive and negative serial dependence is purely due to the changes in environmental statistics *per se*. Other studies also separately used different stimulus features to demonstrate attractive biases for stable attributes such as the identity or gender of a face and repulsive biases for changeable features such as the facial expressions^2,23^. In contrast with these previous studies in which various experimental manipulations interfered with ongoing sequential process, current study only manipulated the environmental statistics with all the other experimental variables kept constant and clearly demonstrated an adaptive switching dynamics between attractive and repulsive bias while observers performed the exactly same perceptual decision task of estimating the mean orientation (Figure 3 & 5). In this sense, our new finding of variance-adaptation effect on serial dependence in ensemble representation cannot be explained by prior studies.

Our results also shed new light on the role of variance adaptation in modulating the interplay between integration and segregation of visual events over time. Our findings demonstrate that, in the same observer, the direction of serial dependence by low-to-low-, high-to-high-, and high-to-low-variance stimuli was positive whereas the direction of serial dependence by low-to-high-variance stimuli was negative (Figure 3 & 5). The dominance of positive serial dependence in most environmental changes implies that humans may have developed a natural prior about the ordered external environment through their experience. Only when the environment gets more disordered as in low-to-high-variance stimuli, human observers adaptively break away from their tendency toward previous information. On the other hand, the opposite directions of serial dependence by low-to-high- and high-to-low-variance stimuli (Figure 3 & 5) indicate that adaptations are stronger after low-variance stimuli, and their repulsive influence increases in relation to future high-variance stimuli. Furthermore, the fact that the direction of bias depends on the temporal increase and decrease of ensemble variance (Figure 3 & 5) and the changes in ensemble variance after the longer history of the same ensemble variance further increase the half-amplitudes of the two opposite biases (Figure 4) suggests that variance adaptation plays a critical role in determining the relative predominance of integration or segregation process. These are in contrast with previous study in which strong adaptation after high-contrast stimuli is rather suppressed by perceptual adjustment process and hence strong signals exert a large attractive bias effect on future low-contrast weak signals^4^. It is possible that current low-contrast weak signals are less weighted than previous high-contrast strong signals during serially dependent perceptual decisions. On the other hand, low-to-high-variance stimuli are likely to signal a drastic context shift, which initiates adaptation processes to segregate representations and reduce the impact of previous information.

Finally, we provide a unified account of the observed opposite directions of serial dependence based on an adapted Bayesian observer model^13,14^. In the present study, a conventional Bayesian observer model is constrained by assuming that 1) adaptation unevenly alters the signal-to-noise ratio around the mean orientation of each Gabor array adaptor and 2) a Bayesian integration is weighted with the most recent stimulus rather than a mixture of past stimuli in a constantly changing environment^6,19–21^. Despite the limitation that this model is not a full sequential Bayesian update model, it does capture the observed adaptive switching behavior between attractive and repulsive recency bias depending on the direction of change in ensemble variance (Figure 6). Along a similar line, the recently proposed hierarchical Two-process model formulated serial dependence as a result of the interaction between low-level adaptation in a sensory unit and weight changes in a decision unit^4^. This Two-process model also provides an account of the coexistence of positive and negative dependencies as in the adapted Bayesian observer model of the present study. In fact, as Chelazzi et al. (2018) pointed out, the Two-process model can be easily accommodated into the predictive coding framework by substituting decisional weights with probability distributions (priors) and low-level population activity with sensory evidence (likelihoods). Thus, both models suggest that serial dependence arises from an integrative process that carries over information from one trial to the next, biasing stimulus appearance. These two models, however, show a clear difference in how the direction of serial dependence is determined. Whereas the plasticity at the level of decisional weights is a critical factor in the Two-process model, the asymmetric change in the likelihood due to the low-level variance adaptation determines the direction of serial dependence in the adapted Bayesian observer model. Therefore, the Two-process model predicts that decisional weight update triggered by the requirement and execution of a behavioral response compensates for trial-by-trial adaptation at the lower level, leading to positive serial dependence. Only when sequential decisions are absent, negative serial dependence appears in this model. However, the adapted Bayesian observer model shows that temporal segregation and repulsive bias do prevail even in the presence of sequential decisions (Figure 6D). These results are in line with the suggestions that serial dependence originates from optimal perceptual strategies based on adaptive coding and temporal integration^24^. Although optimality cannot be guaranteed due to the lack of a full sequential integration modeling in the present study, our results illustrate that the Bayesian observer model constrained by sensory adaptation provides a unified account of serial dependence.

## Conclusion

To conclude, in the present work we provide a unified account of the context-dependent flexibility of serial dependence based on a Bayesian estimation framework constrained by sensory adaptation that allocates more resources to the representation of the parameter values in the vicinity of the adaptor^13,14^. By manipulating only the environmental statistics without changing any of the mean orientation estimation task structure, we clearly demonstrate that the spatiotemporal variability initiates control process to determine which of the two separate and opposing integration and differentiation process dominates. Furthermore, model fitting of behavioral data indicates that the interaction between the process of maintaining visual stability and the process of detecting visual change is automatically and adaptively adjusted mainly through sensory adaptation modulating the encoding efficiency, shaping our visual experience under constantly changing environment. Our study implies that humans have an *a priori* prior about an orderly and structured world rather than a disordered and structureless world, which promotes the temporal integration to a default process in most situations. Only when this fundamental prior belief about a regular world is strongly violated as in low-to-high-variance stimuli (Figure 3-5), the segregation process supporting sensitivity to the environmental stimulus change becomes dominant enough to override the default integration process supporting stability.

## Material and Method

### Observers

A total of 27 observers participated in the experiment; the data from all observers were analyzed. 9 observers (6 males, 3 females) participated in repetitive session. 9 observers (6 males, 3 females) participated in random session. 9 observers (5 males, 4 females) participated in alternating session. All observers had normal or corrected-to-normal visual acuity, gave informed written consent to participate as paid volunteers, and were tested individually in a dark room. The Institutional Review Board of Sungkyunkwan University and the Institutional Review Board of the Korea National Institute for Bioethics Policy approved the study. All participants were naïve to the purpose of the experiment.

### Apparatus

The experiments were conducted in a dark room using the Psychophysics Toolbox along with custom scripts written for MATLAB^25,26^ on a Mac mini. The 22” CRT (ViewSonic PF817) monitor was set to a 100 Hz refresh rate and a resolution of 800 × 600 pixels. The luminance calibration was performed using a calibrated photocell, and monitor gamma tables were adjusted to ensure response linearity and a constant mean luminance of 22.6 cd/m^2^. The observer’s viewing distance from the monitor was 60 cm. Observers used a mouse for all responses (left or right mouse move for adjusting the response bar, and left mouse button click for making the final response), and a spacebar for initiating each block.

### Stimulus

We presented the four-by-four array of sixteen tilted Gabor patches at 12° eccentricity on the right or left visual field. Eccentricity was the visual angle distance from the fixation to the center of the Gabor array. The Gabor patches were located within 10°-by-10° square, and inter-spacing between Gabor patches was 2.5°. The Gabor patches had a peak contrast of 80% Michelson, a spatial frequency of 2.5 cycles per degree, and a 0.5° standard deviation Gaussian contrast envelope. On each trial, orientations of Gabor patches were uniformly distributed with standard deviation either 5° (Low-variance: L) or 20° (High-variance: H) and a mean that was randomly selected from nine possible angles (−40° to 40° in steps of 10°), which differed from the mean orientation of the previous trial. The Gabor array was presented for 1 s and followed by a noise mask of randomly shuffled black and white pixels presented for 1 s at the same location (Figure 1A). The size of the noise mask was same as the Gabor array. After a 300 ms delay, a response bar (width: 0.2°, length: 5°, color: white) appeared at the location of the central fixation point that was a 0.2°-diameter white dot. Observers were asked to adjust the response bar to match the perceived ensemble orientation of the Gabor array using the left/right mouse drags. The starting orientation of the response bar was randomized on each trial. Observers were allowed to take as much time as necessary to respond and pressed the left mouse click to confirm the chosen bar orientation. After a 500 ms delay, the next trial started. Observers were instructed to maintain fixation for the duration of every single trial.

### Procedure and design

There were three different experimental sessions depending on how we manipulated the transition probabilities between the low-variance (L) array and the high-variance (H) array of the consecutive trials (Figure 1B): repetitive (transition probability = 0), random (transition probability = 0.5) and alternating (transition probability = 1) sessions. In repetitive session, either the L or H array was repeatedly presented over trials. In random session, the L and H arrays were randomly presented over trials. In alternating session, the L and H arrays were alternately presented over trials.

Observers performed each of the above three different sessions twice. Each session was composed of 4 blocks of 90 trials (i.e., 360 trials per session). We gave them a break in between blocks and let them freely choose when to start the next block. Observers were not informed of the sequential array statistics of their participating sessions except for the alternating session. Before the alternating session, we informed observers that the L and the H trials would alternate by running several practice trials.

### Data Analysis

Observers’ errors on the adjustment task were calculated as the angle difference between the response bar orientation and the mean orientation of the current Gabor array (current bar orientation – current mean; y-axis on Figure 2A, 3A, 4A, 4C, 5A, 6D). For each observer, trials were treated as outliers if errors exceeded ±3 standard deviations from the mean (on average, 1% of the trials excluded) or no response was made during the response bar adjustment period (on average, 2% of the trials discarded). The first trial of each block was also excluded because there was no previous trial. After outliers were excluded, the response errors were corrected to remove each observer’s general clockwise or counter-clockwise response bias. In order to determine whether an observer’s perception of current mean orientation was influenced by the previous mean orientation, the relative orientation of the previous trial was computed as the angle difference between the previous mean orientation and the current mean orientation (previous mean – current mean; x-axis on Figure 2A, 3A, 4A, 4C, 5A, 6D).

#### Serial dependence fit

In order to quantify the influence of the previous trial on current perceived mean orientation in all three experimental sessions, we fit a simplified derivative of Gaussian (DoG) to each observer’s data using the following equation:

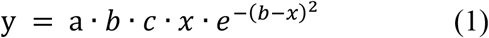

where the parameter ***y*** is the response error on each trial, ***x*** is the relative orientation of the previous trial, ***a*** is half the peak-to-trough amplitude of the DoG, ***b*** scales the width of DoG, and ***c*** is a constant 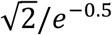, which scales the curve to make the ***a*** parameter equal to the peak amplitude. We fit the DoG using Nelder-Mead method for the nonlinear minimization of the residual sum of squares. As a measure of serial dependence, we used the half amplitude of the DoG curve (paramenter ***a***; Figure 2B, 3B, 4B, 4D, 5B, 7E).

#### Consecutive trial combinations

The serial dependence amplitude was computed for each possible combination of the sequential array statistics between the previous trials and the present trials. In repetitive session, there were only two kinds of consecutive trial combinations of LL (rep L) and HH (rep H). In random session, there were four different consecutive trial combinations of LL, HH, HL, and LH. To examine the effect of exposure duration to a stationary environment on current perceived mean orientation, we also computed serial dependence amplitudes by the consecutive high-to-low-variance and low-to-high-variance trials when the high-variance and low-variance trials were repeated twice and three times before the consecutive trials HHL and HHHL (LLH and LLLH). We performed a linear trend analysis on these one-, two- and three-back amplitudes to test if the serial dependence amplitude gradually decreases or increases as the adaptation to the same spatial variance gets longer. In alternating session, there were two kinds of consecutive trial combinations of LH (alt LH) and HL (alt HL).

#### Mean bootstrapped half amplitude

To be extra-careful on the possibility that few outliers may distort the general serial dependence fit, we used bootstrapping approach to calculate the half amplitudes. For each consecutive trial combination and observer, DoG was fitted with replacement of 5000 times to generate a bootstrapped half-amplitude distribution. The average of half-amplitude distribution was taken as individual’s serial dependence and used for further analysis.

#### Half-amplitude permutation analysis

Significance testsing of mean bootstrapped half-amplitude was conducted with a permutation test. For each consecutive trial combination, we randomly resampled and shuffled the trials’ relative orientation, then recalculated the DoG fit of the shuffled data. This was ran for 10,000 iterations in order to generated a null distribution of half amplitude values. P-values were the proportion of group null distribution that was greater or smaller to the observed half amplitude. Current permutation approach selectively broke the matching relationship between the relative orientation and error, and left the relationship between the stimulus orientation and error untouched. Therefore, the permutated null distribution automatically includes unwanted serial dependence caused by other factors such as the oblique effect. By applying permutation analysis, undesired effect on serial dependence can be nullified.

#### Combination permutation analysis

To test whether the half-amplitude of certain combinations is distinctive from the others, permutation on half-amplitude of combinations were performed. For each iteration, we shuffled the mean bootstrapped half-amplitude of all observers and combinations, then obtained a permutated half-amplitude averaged across observers. This procedure was repeated for 10,000 times, which result in the null distribution. P-values were the proportion of group null distribution that was greater or smaller to the observed half amplitude, and alpha criterion was Bonferroni-corrected (i.e. divided by 4 in random session).

### Modeling

To measure the effect of variance adaptation on ensemble serial dependence, a model in Bayesian perceptual inference framework was built for each combination of two consecutive trials in random session (i.e. LL, HH, HL, and LH) as the following formula:

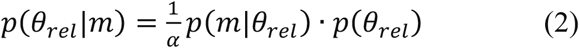

where *θ*_*rel*_ denotes the relative orientation between the previous and the present mean orientation, and m as each measurement of the encoder with normalization factor α.

Of the profile of likelihood function of each measurement, normal distribution on the relative orientation domain was assumed;

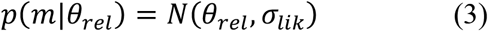

where *σ*_*lik*_ denotes the standard deviation of each measurement. Critically, unevenly higher encoding reliability around the adapted orientation in relative orientation domain was supposed, which is the assumption adopted form the previous studies [3–5].

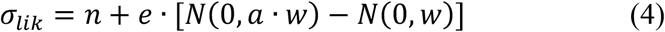

*σ*_*lik*_ was composed of constant sensory noise, n, encoding effect size, e, and encoding effect width, w, with constant a which was two. The zero degree of the relative orientation denotes the orientation that the previous stimulus presented. Sensory noise, n, regulates overall noise, and fitted separately for the cases of current trial presents L and H. That is, LH and HH trials (HL and LL trials) had the same sensory noise parameter n, since sensory noise supposed to purely depend on what is currently on visual. Encoding effect meant that diminishing effect on sensory noise in the vicinity of the adaptor. It was comprised of the summation of two normal distributions; one reduces sensory noise immediate nearby the adapted orientation, and the other increases sensory noise beyond the very close adaptor (Figure 6a). It was under the assumption that the additional resource to sharpen encoding around the adaptor was required to be compensated. The encoding effect was modulated by two parameters. The first parameter was encoding effect size, e, which adjusts the relatively reduced amount of sensory noise; e being equal to one means the sensory noise is decreased roughly to half near the adapted orientation, compared to that of distant orientation. The second parameter was encoding effect width, ***w***, which controls the standard deviation that the encoding effect influences. With varying encoding effect size, e, and effect width, w, the likelihood function may skew (Figure 6b).

Of the profile of prior probability distribution, Gaussian mixture prior was assumed (Figure 6c), which the assumption adopted from the previous study [6]. The prior was composed of maximum value between the normal distribution and flat prior. The flatter prior utilized meant the perception valued more on sensory evidence, vice versa. The prior distribution was expressed as:

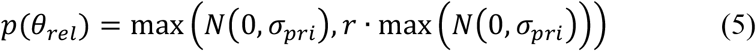

The prior distribution was controlled by two parameters; the standard deviation of the normal distribution, *σ*_*pri*_, modulated the width prior impact on, and flat prior ratio, r, adjusted the weight between flat and Gaussian prior. If the flat prior ratio, r, was one, no prior knowledge was used for perceptual inference.

Finally, the mean of posterior probability distribution was taken as the optimal estimate:

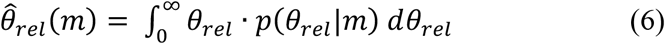

We fit the average response error data to above model for each of four different consecutive trial combination. In order to observe group-wise fitness, the model was fitted to data regardless of observers (Figure 6d), and to test the significance, the model was fitted to individual’s data (Figure 6e, f), and tested under permutation t-test. The permutation was performed under null distribution with shuffling 10,000 times the combinations and observers, and p-values were Bonferroni corrected.

